# Amino Acid Composition and Charge Based Prediction of Antisepsis Peptides by Random Forest Machine Learning Algorithm

**DOI:** 10.1101/2021.09.26.461860

**Authors:** Aayushi Rathore, Anu Saini, Navjot Kaur, Aparna Singh, Ojasvi Dutta, Mrinal Bamhotra, Avneet Saini, Sandeep Saini

## Abstract

Sepsis is a severe infectious disease with high mortality, and it occurs when chemicals released in the bloodstream to fight an infection trigger inflammation throughout the body and it can cause a cascade of changes that damage multiple organ systems, leading them to fail, even resulting in death. In order to reduce the possibility of sepsis or infection antiseptics are used and process is known as antisepsis. Antiseptic peptides (ASPs) show properties similar to antigram-negative peptides, antigram-positive peptides and many more. Machine learning algorithms are useful in screening and identification of therapeutic peptides and thus provide initial filters or built confidence before using time consuming and laborious experimental approaches. In this study, various machine learning algorithms like Support Vector Machine (SVM), Random Forest (RF), K-Nearest Neighbour (KNN) and Logistic Regression (LR) were evaluated for prediction of ASPs. Moreover, the characteristics physicochemical features of ASPs were also explored to use them in machine learning. Both manual and automatic feature selection methodology was employed to achieve best performance of machine learning algorithms. A 5-fold cross validation and independent data set validation proved RF as the best model for prediction of ASPs. Our RF model showed an accuracy of 97%, Matthew’s Correlation Coefficient (MCC) of 0.93, which are indication of a robust and good model. To our knowledge this is the first attempt to build a machine learning classifier for prediction of ASPs.

## 1. INTRODUCTION

Over the last few decades, sepsis has become a leading cause of deaths in non-coronary intensive care units [1], and is still an ill-defined disease characterized by complex and time-dependent pathophysiological processes [2]. In the 1950s, gram-positive bacteria were the primary cause of sepsis, but after the introduction of antibiotics, gram-negative bacteria became the predominant cause of sepsis from the 1960s to the 1980s [3]. In clinical terms, sepsis is an inflammatory dys-regulation exhibited after bacterial infection; and is categorized into sepsis: an inflammatory immune response triggered by an infection [4, 5], severe sepsis: causing poor organ function or blood flow, and septic shock: low blood pressure due to sepsis that does not improve after fluid replacement [6]. The mortality rate associated with sepsis is as high as 30%, as high as 50% from severe sepsis, and up to 80% from septic shock, [7] moreover sepsis affected about 49 million people in 2017, with 11 million deaths worldwide [8], and contributes over 6% of total deaths in the United States. As we are exposed to potentially dangerous pathogens daily, techniques must be applied to eliminate contamination present on objects and the skin employing sterilization and disinfection.

The process of prevention of infection by inhibiting or arresting the growth and multiplication of germs is known as antisepsis; the antimicrobial products used for this purpose are known as antiseptics, which are applied to living tissue or skin to reduce the possibility of infection, sepsis, or putrefaction; certainly, the peptides which kill the gram-negative and gram-positive bacteria are known as antiseptic Peptides [9]. These peptides also show properties similar to antigram-negative peptides, antigram-positive peptides, anti HIV peptides, antiparasitic peptides, antimalarial peptides, hemolytic peptides, insecticidal and hence demonstrate potential as novel therapeutic agents [9].

As peptides are recognized for being highly tuneable molecules, they can be tailored to achieve desirable biocompatibility and biodegradability with simultaneously selective and potent therapeutic effects, thus there is an increased interest in peptides in clinical therapeutics [10]. In general, peptides are selective and efficacious signaling molecules that bind to specific cell surface receptors, such as G protein-coupled receptors (GPCRs) or ion channels, where they trigger intracellular effects; and because of their attractive pharmacological profile and intrinsic properties, they represent an excellent starting point for the design of novel therapeutics and their specificity has been seen to translate into excellent safety, tolerability, and efficacy profiles in humans. This aspect might also be the primary differentiating factor of peptides compared with traditional small molecules. Furthermore, peptide therapeutics is typically associated with lower production complexity compared with protein-based biopharmaceuticals and, therefore, the production costs are also lower. Thus, peptides are in the sweet spot between small molecules and biopharmaceuticals; but they need to be classified from other peptides [10].

Antiseptic peptides can be classified from other peptides based on the following two methods: Wet Lab and Model Building. Amongst the two, the Model Building approach is mostly used because identifying suitable therapeutic peptides via traditional experimental approaches involves screening of peptide libraries, or examining whole proteins using overlapping window patterns in regions of peptide chains and then assessing each part for its activity, which is a time consuming, costly, and laborious process [11]. Various machine learning algorithms are been developed to rapidly and efficiently predict the utility of therapeutic peptides and offer an array of tools that can accelerate and enhance decision making and discovery for well-defined queries with ample and sophisticated data quality. The use of these tools in the development of peptide-based therapeutics is already revolutionizing protein research by unravelling their novel therapeutic peptide functions [11].

In this work, we evaluated the potential of different physiochemical properties of ASPs to build first machine learning model for antisepsis peptides.

## 2. MATERIAL AND METHODS

### 2.1 Data collection and processing

Data collection and processing is the initial step involved in the construction of machine learning based models. Along with the various databases, extensive literature searches have been performed, the dataset obtained includes experimentally verified positive samples. A total of 76 entries of anti-sepsis peptides were retrieved out of which 75 were from the APD3 database and 1 literature.

The negative dataset can consists of the following peptides: (a) random peptides from UniProt; (b) random shuffling of positive samples; and (c) peptides with different functions rather than the desired function. This approach is based on the hypothesis stating that the chances of a negative sample presenting as a false negative are minimal (Shaherin Basith et al, 2019). The final negative set that is used in our study consists of 197 entries of randomly retrieved peptides from UniProt.

After the generation of the positive and the negative dataset, the removal of the redundant peptides is done using CD-HIT webserver as the redundant samples may overestimate the prediction accuracy of the model. For machine learning model development 80% of the original data is used as training set and the rest 20% as independent data set for testing of the developed models.

### 2.2 Feature Extraction

Features acts as a discriminatory attribute for distinguishing between the anti-sepsis and non anti sepsis peptides, to identify the anti-sepsis peptides by means of sequence alone, we have to convert the plain amino acid sequence into numerical descriptors characterizing different properties of peptides. For the binary classification amino acid composition (AAC) and dipeptide composition (DPC) were computed using iFeature, a webserver for peptide features extraction. The values of pseudo amino acid composition (PseAAC) were obtained using webserver Pfeature. Other physiochemical properties like length of amino acids, Boman index, hydrophobicity, net charge, molecular weight and isoelectric point (pI) were retrieved from the APD3 database along with positive dataset of ASPs.

### 2.3 Feature Selection

Feature selection is a method used to select a subset of relevant features for model building. In this study it has been done via two methods; the first one is by using python based package RFE, recursive feature elimination and other is manual selection. Out of total 447 features, RFE selected 8 best features for model building. In case of manual selection, a number of combinations have been made and checked against different machine learning algorithms.

Best model was trained on only 2 features were selected for further evaluation, which includes net charge and amino acid composition.

### 2.4 Construction of Machine Learning Based Models

Machine learning explores the construction and evaluation of algorithms that can be used for analysis, classification and prediction based on models learned from existing data (Maryam Mousavizadegan et al, 2016). Python sci-kit leran machine learning package is used for model development and validation. A number of attempts have been made for the construction of machine learning based tool using the various classifiers; which includes SVM (Support Vector Machine), RF (Random Forest), KNN (K-Nearest Neighbours) and LG (Logistic Regression). Out of these, the accuracy of the RF classifier was maximum. Thus, considered the best approach for the prediction of anti sepsis peptides.

The various algorithms or classifiers that were applied to the ASPs dataset include:

I. Random Forests is an ensemble learning algorithm which operates by setting up a combination of identically distributed decision trees in a way that each tree depends on the value of a random vector sampled independently. Each tree plays an equal part in voting for the most popular class .Random forests have been used for classification of various biological data such as microarray gene expression data, proteomics data, protein-protein interaction data and sequence analysis (Maryam Mousavizadegan et al, 2016).
II. The objective of support vector machines (SVMs) in binary classification problems is to obtain the best hyper plane that separates the two classes, thus minimizing the error. Te hyper plane is defined through support vectors. Since most real problems do not have a linear relationship, the SVM algorithm offers the possibility of calculating a kernel function to map the data in a greater number of dimensions, making it possible to linearly separate the data.
III. The K-nearest neighbor (k-NN) algorithm is a technique based on cluster theory. It is a very basic algorithm, but it has been reported to yield excellent results for classification. It is based on the fact that a new observation that is particularly close to an observation within the learning set should have a great weight in the decision and, conversely, an observation that is at a farther distance will have a much smaller weigh. A very high k can cause over-training of the model; the decision was made to maintain intermediate levels. The range of values used was from 1 to 5.
IV. Logistic regressions are popular classification algorithms in machine learning problems when the response variable is categorical. Te logistic regression algorithm represents the class-conditional probabilities through a linear function of the predictors (Jose Linares Blanco et al,2018).

### 2.5 Cross Validation Measure

We used repeated k-fold cross validation to evaluate the performance of the predictive model used in this study. In the K-fold cross-validation, the whole data set is randomly split into K sets, out of which one set (test) is tested by a model developed on the remaining K-1 sets (training). This process is then iterated for a number of times. In our study we used five-fold cross validation.

### 2.6 Performance Evaluation

ROC (Receiver Operating Characteristic) and CAP (Cumulative Accuracy Profile) curve were used for the model analysis. We used sensitivity, specificity, accuracy, precision and MCC to evaluate the performance of the model. They can be calculated using the following formulae:

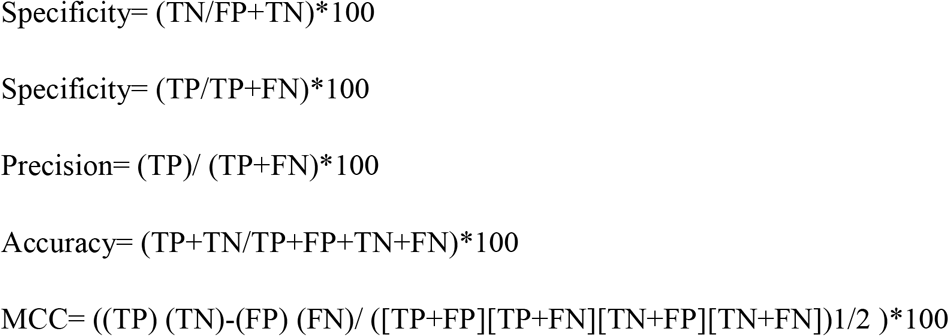

Where TP and TN are correctly predicted True positive and True negative examples, respectively. Similarly, FP and FN are wrongly predicted False positive and False negative examples, respectively, and MCC is the Mathew’s correlation coefficient.

## 3. RESULTS AND DISCUSSION

### 3.1 Exploratory Data Analysis

Exploratory data analysis refers to the critical process of performing initial investigations on data so as to discover patterns, to spot anomalies, to test hypothesis and to check assumptions with the help of summary statistics and graphical representations. Exploratory data analysis is an approach to analyse data sets to summarize their main characteristics, often with visual methods.

#### a.) Compositional Data Analysis

Before we develop in silico models for prediction, it is important to analyze Antisepsis peptides (ASPs) to understand their nature. As all peptides are made of 20 types of residues, it is important to examine the frequency of each type of residues in ASPs. (Piyush et al, 2018). Thus, we computed and compared the amino acid composition of ASPs and non-ASPs of our dataset. The average AAC of the two groups (ASPs and non-ASPs) are displayed using a bar Graph, made using matplotlib library from python as shown in **Fig 1**. The analysis showed that certain residues like A, C, F, G, K, L, P, R and V, are more abundant or frequent in ASPs whereas non-ASPs are dominated by residues like A, F, G, I, K, L, T and V. Presence of residues like C, K, R makes ASPs positively charged and cationic in nature.

**Figure 1:**
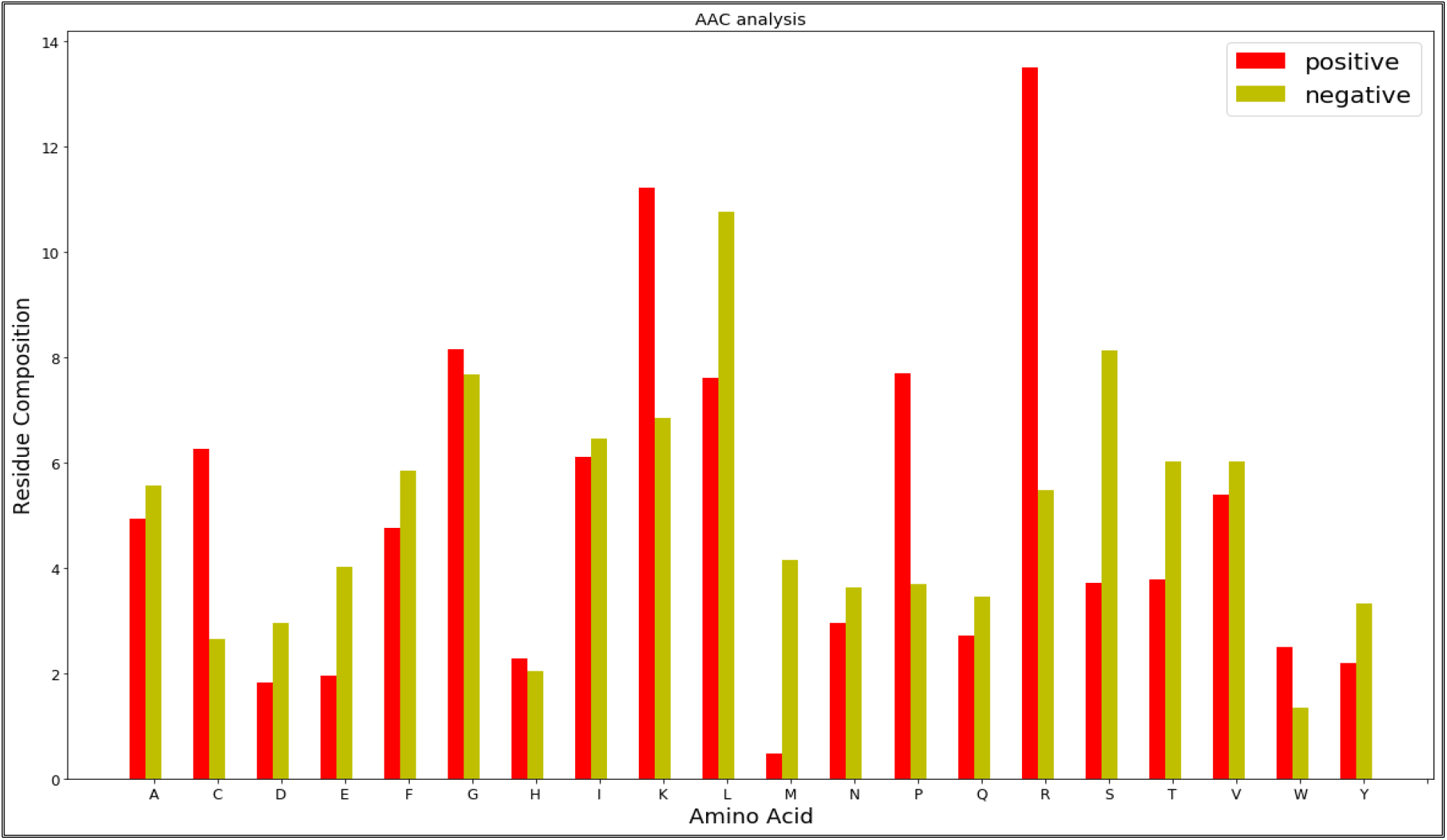
Amino Acid Composition (AAC) Analysis of Positive & Negative Data Set.

#### b.) Physiochemical Property Analysis

Along with the compositional features physiochemical properties such as charge, pI, length, and hydrophobicity etc. are also important and in order to find the importance of each physiochemical property box plots were drawn using python matplotlib library as shown in **Fig 2**.

**Figure 2:**
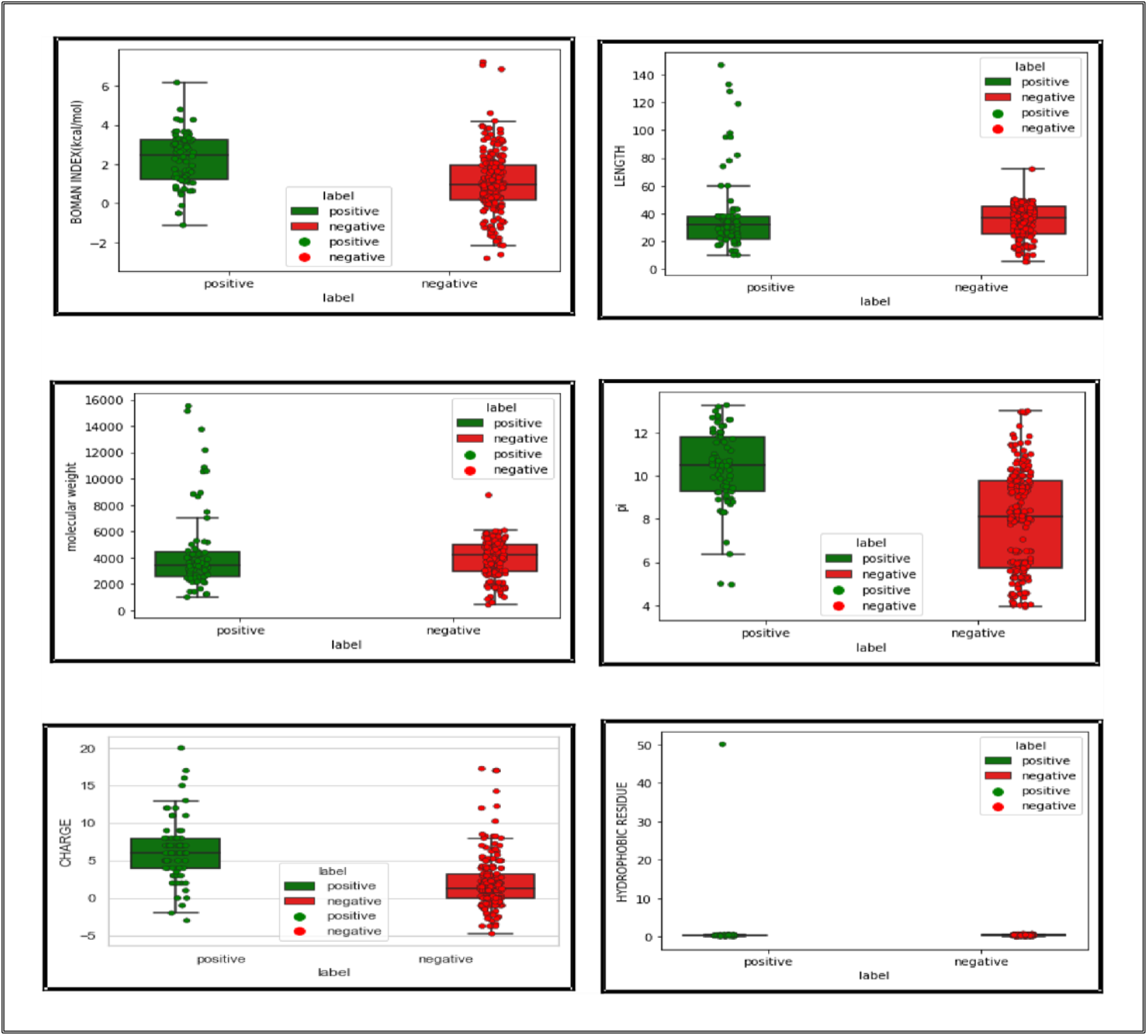
Physiochemical Property Analysis of Positive & Negative Data Set.

### 3.2 Effect of ASPs and non-ASPs peptide data ratio

As experimental non-ASP data are scarce, a conventional way to produce negative samples is selection of random sequences from a protein database following certain criteria or by shuffling the residues of the ASP peptides. This approach enables generation of a large amount of negative data. For training a classifier, one would usually select a positive/negative (P:N) data ratio of 1:1. By contrast, in a study by Li et al on rebalancing data ratio techniques for inherently imbalanced medical data, they showed that in some cases, optimal classification accuracy can be achieved with a slight imbalance of the data distributions (Bhadra et al, 2018). Therefore in our study we created two datasets with ratio 1:1 and 1:2 and evaluated the predictive performances of each RF classifier with 21 features by five-fold cross validation and results are demonstrated in **Table 1**.

**Table 1:** Predictive performances with different data ratios of positive and negative dataset ratios.

| **Data ratio** | **Accuracy** | **Sensitivity** | **Specificity** | **MCC** | **Precision** |
| --- | --- | --- | --- | --- | --- |
| 1:1 | 0.16 | 0.3 | 0.07 | -0.65 | 0.18 |
| 1:2 | **0.97** | 1.00 | 0.90 | 0.93 | **0.97** |

### 3.3 Feature Extraction and Analysis

Features of both the peptide groups (positive and negative) were calculated which made a total of 447 features including AAC, DPC, PseAAC, pI, molecular weight, Length, hydrophobicity, Boman Index and net Charge using iFeature, Pfeature and APD3 Calculation and Prediction tool. After performing manual as well as python package called RFE (Recursive Feature Elimination) feature selection (Atanaki et al, 2020), total of 2 and 8 features were selected whose performance matrices are given in **Table 2**. As per the results with RF model manual feature selection was used in the final model which included AAC and net Charge as features.

**Table 2:** Predictive performance with different feature extraction methods.

| **Feature selection Method** | *Accuracy* | *Sensitivity* | *Specificity* | *MCC* | *Precision* |
| --- | --- | --- | --- | --- | --- |
| **RFE (8 features)** | 0.93 | 0.93 | 0.90 | 0.81 | 0.96 |
| **Manual (2 features)** | 0.97 | 1.00 | 0.90 | 0.93 | **0.97** |

### 3.4 Model Evaluation with different classifiers

Different classifiers such as SVM, KNN, RF, LR and NB were used in order to get the best model. **Table 3** shows the performance of the models created using different classifiers. Also an ROC curve i.e. receiver operating characteristic curve that plots TP rate (sensitivity) and FP rate (specificity) and a CAP curve i.e. cumulative accuracy profile curve that analyse to effectively identify all data points of a given class using minimum number of tries. **Figure 3** and **Figure 4** respectively shows the curves for evaluation of the classifiers.

**Table 3:**
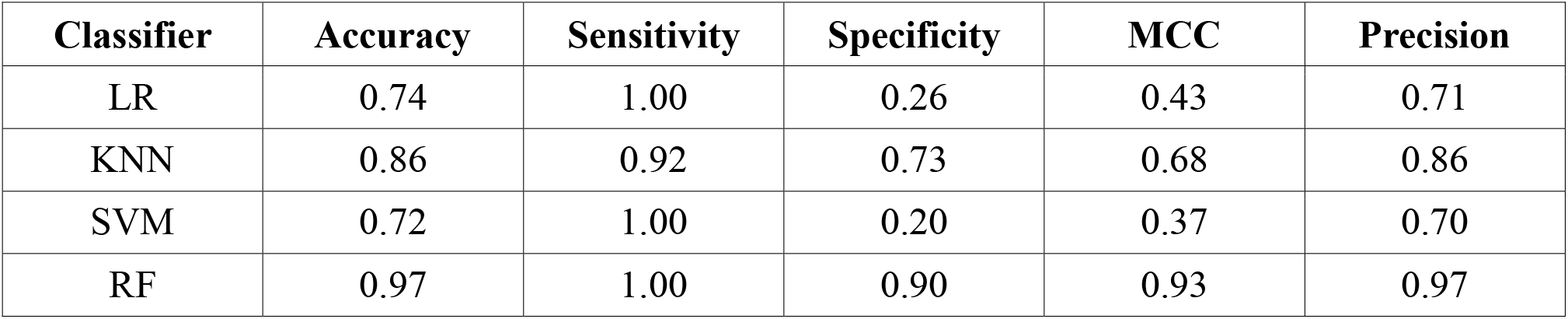
Predictive performance with different machine classifiers or algorithms.

| Classifier | Accuracy | Sensitivity | Specificity | MCC | Precision |
| --- | --- | --- | --- | --- | --- |
| LR | 0.74 | 1.00 | 0.26 | 0.43 | 0.71 |
| KNN | 0.86 | 0.92 | 0.73 | 0.68 | 0.86 |
| SVM | 0.72 | 1.00 | 0.20 | 0.37 | 0.70 |
| RF | 0.97 | 1.00 | 0.90 | 0.93 | 0.97 |

**Figure 3:**
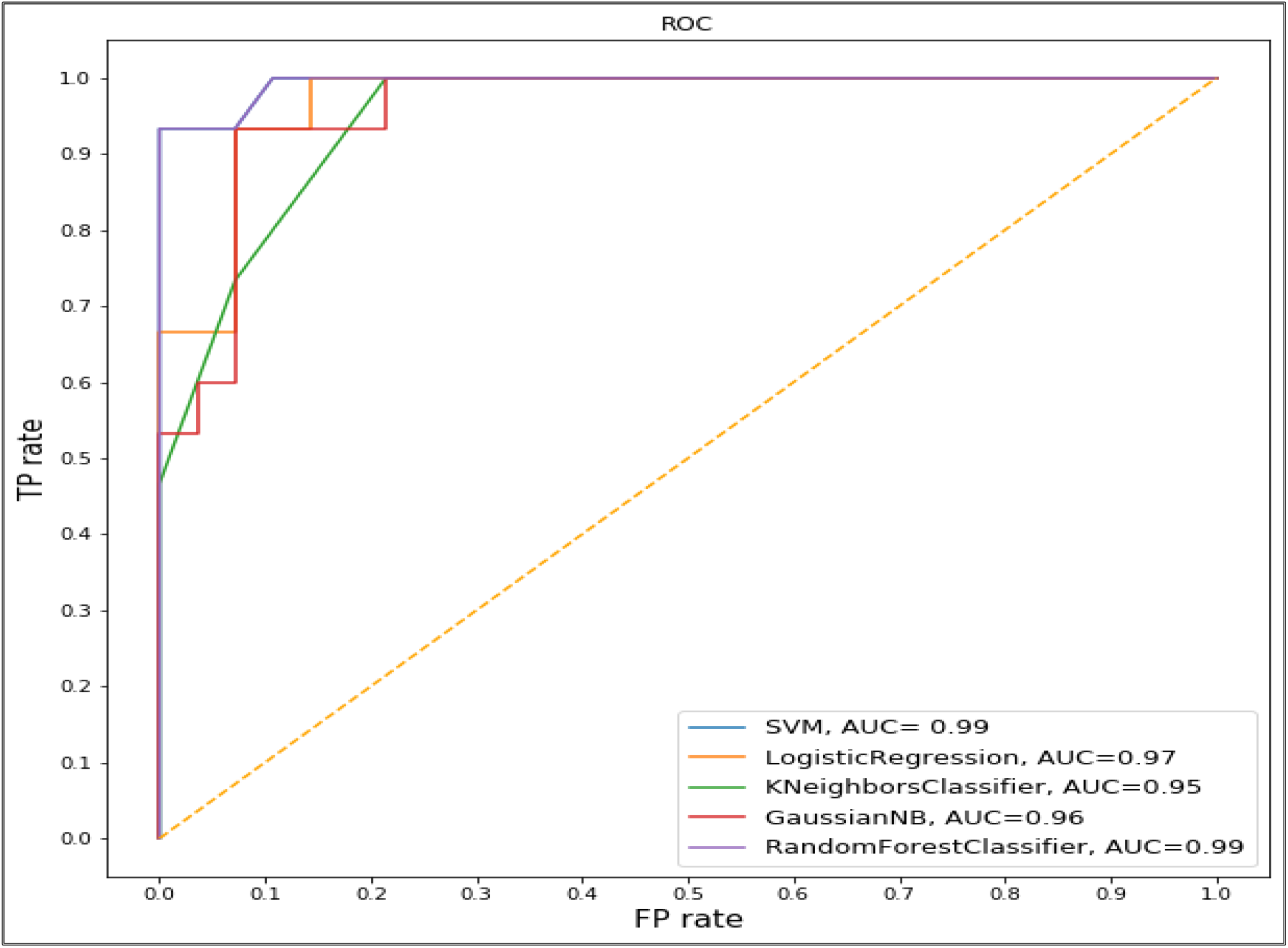
ROC curve showing sensitivity and specificity for classifiers SVM, LR, kNN and RF.

**Figure 4:**
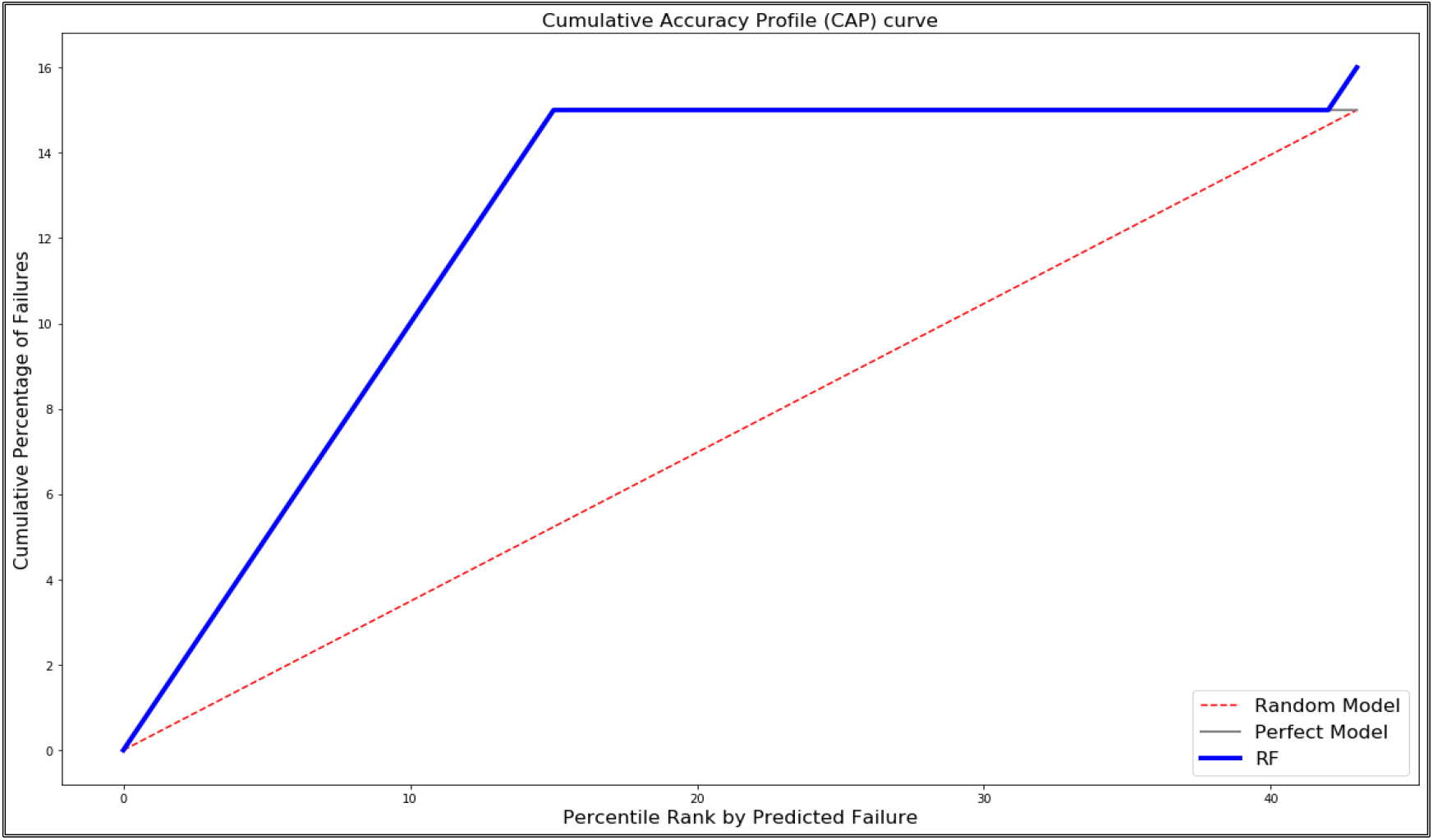
CAP curve shows the percentile rank by predicted failure and cumulative percentage of failures for Random Model, Perfect Model and RF Model (Random Forest).

### 3.5 Performance comparison of independent dataset with existing model

In statistical prediction, cross-validation methods are often used to examine a predictor for its effectiveness in practical application (Schaduangrat et al, 2019). We used five-fold cross-validation along with the independent dataset to evaluate the performance of the predictive model used in our study and finally five-fold cross validation model was selected based on the performances provided in **Table 4**. In the five-fold cross-validation, the whole data set is randomly split into five sets, out of which one set (test) is tested by a model developed on the remaining four sets (training). This process was then iterated 150 times (Atanaki et al, 2020).

**Table 4:**
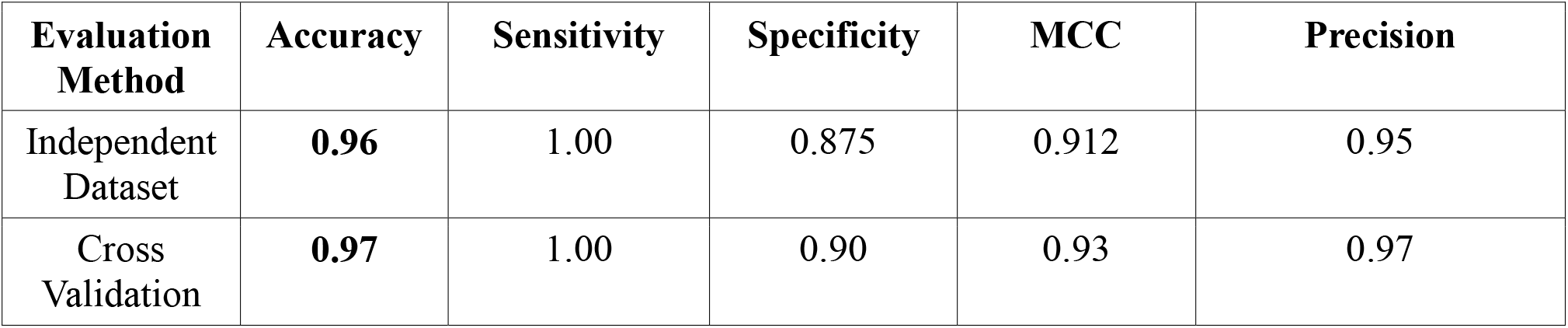
Predictive performance of RF algorithm after independent and cross-validation approaches.

| Evaluation Method | Accuracy | Sensitivity | Specificity | MCC | Precision |
| --- | --- | --- | --- | --- | --- |
| Independent Dataset | **0.96** | 1.00 | 0.875 | 0.912 | 0.95 |
| Cross Validation | **0.97** | 1.00 | 0.90 | 0.93 | 0.97 |

## DISCUSSION

Sepsis is life-threating organ dysfunction condition due to a deregulated host response to infection (Kristin E Rudd et al 2020). It is prevented by inhibiting or arresting the growth and multiplication of germs which is known as Antisepsis. With the help of Antimicrobial products the process of antisepsis is performed. Peptides are selective and constructive signalling molecules which bind to G protein-coupled receptor (GPCRs) or ion channels activating intracellular effects.

In this paper, we have used the machine learning approach for building classification model to differentiate antiseptic peptides from other peptides. Machine learning algorithms such as SVM, RF, KNN, LR and NB were trained and evaluated for performance using test and cross-validation approaches in python machine learning environment, sci-kit learn. ROC and CAP curves analyses assessment of the classifiers were also performed to select the best classifier on the basis of accuracy, sensitivity and specificity.

Positive dataset was created using most of the peptides from APD3 whereas negative dataset peptides were downloaded from Uniprot. Features were extracted to compare the anti-sepsis and non-anti-sepsis peptides. The features like AAC and DPC and other properties like Boman Index, hydrophobicity and net charge etc. were calculated to make total of 447 features representative of peptide datasets.

These extracted features were filtered to 8 with RFE (Random Feature Extraction) which is a python based package and provided us with features like Amino Acid Composition (AAC) and Net Charge which gave the best accuracy but we have taken manually selected features which were 21 in number and gave more accurate model which build using RF (Random Forest). As the model created by RF was 0.97 accurate comparing to the model created by RFE which was 0.93 accurate.

Performance comparison was done of independent dataset with existing model where statistical prediction, cross validation method was used. 10 fold as well as 5 fold cross validation was used along with independent dataset to determine the performance of predictive model where we have taken repeated 5 fold cross validation model based on the performances provided in the results.

## CONCLUSION

Sepsis is a life-threatening illness caused by your body’s response to an infection. The process of prevention of infection by inhibiting or arresting the growth and multiplication of germs is known as Antisepsis. We have taken 76 Antisepsis peptides as Positive dataset and 197 peptides of non-antisepsis activity as Negative Dataset which results the P/N (positive-negative) ratio to be 1:2. Then feature extraction for the both datasets was done by using several tools and total no. of features obtained where 447 out of which 21 were taken by the manual feature selection and 8 were obtained through RFE (Random Feature Extraction) python based package. As per the result model was built using RF classifier (Random Forest) which is a Machine-Learning algorithm with manual feature selection using AAC and Net charge as features which results in 0.97 accuracy of the model, also ROC (Receiver Operating Characteristic Curve) and CAP (Cumulative Accuracy Profile Curve) were obtained for the evaluation of classifiers. Then statistical prediction of independent dataset was done through cross-validation method where repeated five-fold cross-validation model was selected which results in 0.97 accuracy of the model.

## Graphical Abstract

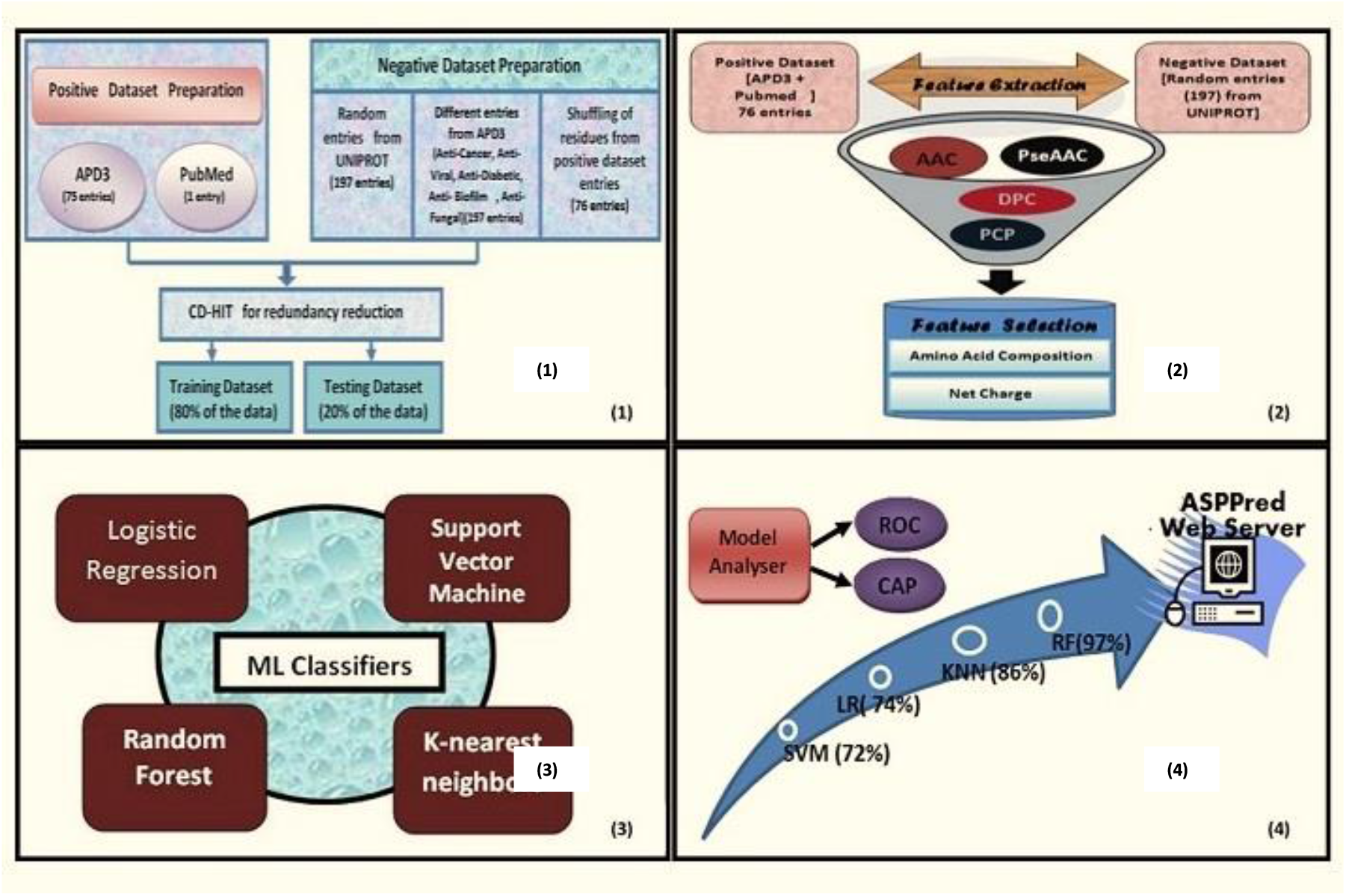

